# Quantitation of tizoxanide in multiple matrices to support cell culture, animal and human research

**DOI:** 10.1101/2021.05.27.445500

**Authors:** Megan Neary, Usman Arshad, Lee Tatham, Henry Pertinez, Helen Box, Rajith KR Rajoli, Anthony Valentijn, Joanne Sharp, Steve P Rannard, Giancarlo A Biagini, Paul Curley, Andrew Owen

**Affiliations:** Department of Pharmacology and Therapeutics, University of Liverpool, Liverpool, L7 3NY, UK; Department of Chemistry, University of Liverpool, Liverpool, L69 3BX, UK; Centre of Excellence in Long-acting Therapeutics (CELT), University of Liverpool, Liverpool, L7 3NY, UK; Centre for Drugs and Diagnostics, Liverpool School of Tropical Medicine, Liverpool L3 5QA, UK

**Keywords:** Tizoxanide, LC-MS/MS, plasma, SARS-CoV-2, COVID-19

## Abstract

Currently nitazoxanide is being assessed as a candidate therapeutic for SARS-CoV-2. Unlike many other candidates being investigated, tizoxanide (the active metabolite of nitazoxanide) plasma concentrations achieve antiviral levels after administration of the approved dose, although higher doses are expected to be needed to maintain these concentrations across the dosing interval in the majority of patients. Here an LC-MS/MS assay is described that has been validated in accordance with Food and Drug Administration (FDA) guidelines. Fundamental parameters have been evaluated, and these included accuracy, precision and sensitivity. The assay was validated for human plasma, mouse plasma and Dulbeccos Modified Eagles Medium (DMEM) containing varying concentrations of Foetal Bovine Serum (FBS). Matrix effects are a well-documented source of concern for chromatographic analysis, with the potential to impact various stages of the analytical process, including suppression or enhancement of ionisation. Therefore, a robustly validated LC-MS/MS analytical method is presented capable of quantifying tizoxanide in multiple matrices with minimal impact of matrix effects. The validated assay presented here was linear from 15.6ng/mL to 1000ng/mL. Accuracy and precision ranged between 102.2% and 113.5%, 100.1% and 105.4%, respectively. The presented assay here has applications in both pre-clinical and clinical research and may be used to facilitate further investigations into the application of nitazoxanide against SARS-CoV-2.

## Introduction

Nitazoxanide (NTZ) is a thiazolide anti-protozoal drug, approved to treat diarrhoea caused by the parasites *Cryptosporidium* and *Giardia* in children and adults [1, 2], and is administered orally as a suspension or tablet twice daily with food [2]. Additional efficacy has been demonstrated both *in vitro* and *in vivo* against a wide range of parasites, bacteria, fungi and viruses [3–9]. This broad-spectrum of activity is in part attributed to NTZs rapid transformation into its deacetylated active metabolite tizoxanide (TIZ) [4]. NTZ has previously been considered for drug repurposing to treat respiratory viral infections [9], including Middle Eastern respiratory syndrome (MERS) [10], with clinical trials having assessed its suitability to treat acute influenza [11, 12].

Currently NTZ is being assessed as a candidate therapeutic for severe acute respiratory syndrome coronavirus 2 (SARS-CoV-2), with 25 trials listed on clinicaltrials.gov [13]. *In vitro* evaluation of the activity of NTZ against SARS-CoV-2 in Vero E6 cells generated an EC_90_ estimated at 4.65 μM [14, 15]. Subsequently, TIZ was additionally confirmed to have antiviral activity with an EC_90_ 3.16 μM under similar assay conditions [16]. Unlike many other candidates being investigated, TIZ plasma concentration achieve antiviral levels after administration of the approved dose [14],although higher doses are expected to be needed to maintain these concentrations across the dosing interval in the majority of patients [17]. Several mechanisms of action for NTZ against SARS-CoV-2 have been proposed and are summarised by Lokhande *et al*. [18].NTZ has previously been demonstrated to inhibit production of pro-inflammatory cytokines and interleukin-6 in peripheral blood mononuclear cells *in vitro* [19], and reduce interleukin-6 plasma levels in mice [20]. It has been hypothesised that this may assist host anti-inflammatory responses during SARS-CoV-2 disease progression and thus lessen the SARS-CoV-2 cytokine storm [9, 18].

Research into the suitability of NTZ as a treatment for SARS-CoV-2 is currently ongoing and includes dose optimisation, tolerability and drug-interaction studies [17, 21]. One uncertainty relating to the putative utility of NTZ is that TIZ is highly protein bound in plasma of humans [22]. However, whether this binding is high affinity / low capacity or low affinity / high capacity is currently unknown and could have a profound impact upon the probability of success [23, 24]. Methods for quantification of concentrations of NTZ and TIZ in a variety of matrices are required to better understand the exposure-response relationship, *in vitro, in vivo* and in humans. Currently few validated liquid chromatography tandem mass spectrometry (LC-MS/MS) assays are available to quantify NTZ and TIZ in relevant biological matrices. Previous studies include a validated LC-MS method for quantifying NTZ and TIZ in mouse plasma, lung, BAL cells, liver, spleen kidney and heart with a limit of quantification (LOQ) of 0.97 ng/mL [25]. As well as a LC-MS/MS methods validated in goat faeces with a limit of detection of 5ng/g [26]. A high performance liquid chromatography (HPLC) method has been validated for NTZ and TIZ in plasma but is anticipated to lack the sensitivity needed to determine the pharmacokinetics of NTZ and TIZ as a SARS-CoV-2 treatment [27].

Here an LC-MS/MS assay is described that has been validated in accordance with Food and Drug Administration (FDA) guidelines [28]. Fundamental parameters have been evaluated, and these included accuracy, precision and sensitivity. Other validation criteria also included linearity, recovery, reproducibility and stability of TIZ within selected matrices. The assay was validated for human plasma, mouse plasma and Dulbeccos Modified Eagles Medium (DMEM) containing varying concentrations of Foetal Bovine Serum (FBS). Matrix effects are a well-documented source of concern for chromatographic analysis, with the potential to impact various stages of the analytical process [29], including suppression or enhancement of ionisation [28]. Therefore, a robustly validated LC-MS/MS analytical method is presented capable of quantifying TIZ in multiple matrices with minimal impact of matrix effects.

## Methods and Materials

### Materials

TIZ and the internal standard (IS) TIZ-D4 were purchased from Toronto Research Chemicals inc (Toronto, Canada). Drug free mouse plasma and human plasma with lithium heparin were purchased from VWR International (PA, USA). LCMS grade acetonitrile (ACN) was purchased from Fisher Scientific (MA, USA). All other consumables were purchased from Merck Life Science UK LTD (Gillingham, UK) and were of LCMS grade.

### Tuning for Tizoxanide and Internal Standard

A stock solution of 1mg/mL of TIZ or TIZ-D4 was prepared in ACN. From this stock a solution of 10ng/mL was prepared in H2O:methanol (50:50) for tuning. Detection was performed using a SCIEX 6500+ Qtrap (SCIEX MA, USA) operating in negative mode. TIZ or TIZ-D4 was infused at 10μL/min in order to optimise compound-specific parameters (declustering potential, collision energy and collision exit potential) and source specific parameters (curtain gas, ionisation voltage, temperature, nebuliser gas and auxiliary gas).

### Chromatographic Separation

TIZ was resolved using a kinetex C18 column (2.7μM, 2.1×100mm Phenomenex CA, USA) using a 5μL injection. The following multistep gradient was used with H_2_O + 0.1% formic acid (mobile phase A) and ACN + 0.1% formic acid (mobile phase B) at a flow rate of 0.4mL/min: the initial conditions were 5% A and 95% B and held for 1 minute. Mobile phase B was rapidly increased to 80% over 0.5 mins. The gradient was then slowly increased to 95% B till 4 mins. This was held for 0.5 mins when the initial conditions were returned and held for 0.5mins.

### Extraction from mouse plasma, human plasma and foetal bovine serum

The assay described here was developed to quantify TIZ from *in vitro* (DMEM plus 50%, 25%, 12%, 10%, 5% and 2% FBS or 50% and 10% human plasma) and *in vivo* (mouse plasma and human plasma) matrices to support SARS-CoV-2 preclinical research. 100μL of standard quality control (QC) sample or untreated matrix was pipetted into 7mL glass tubes. 500μL of ACN containing IS (20ng/mL) was added to each tube and thoroughly vortexed. Following vortexing, the samples were centrifuged at 3500rpm for 5mins at 4°C. 500μL of supernatant was then transferred to fresh 7mL tubes and the samples were evaporated to dryness under a gentle stream of nitrogen. All samples were reconstituted in 100μL of reconstitution buffer (30% H_2_O, 49% ACN and 21% DMF). 50μL was the transferred to chromatography vials and analysed.

### Linearity

Calibrators were prepared by spiking untreated matrix with TIZ followed by serial dilution, ranging from 15.6ng/mL to 1000ng/mL. Linearity was assessed over 3 independent runs. Acceptance criteria were as follows; deviation of interpolated standards from stated concentrations were set at 15%, excluding the lower limit of quantification (LLOQ) where deviation was set at no more than 20%. Acceptable R^2^ was set at >0.99.

### Recovery

Recovery was assessed at 3 concentrations (40ng/mL, 400ng/mL and 800ng/mL) covering the dynamic range of the assay. The peak area of extracted samples was compared to extracted blank matrix spiked with the expected concentrations.

### Matrix Effects

Matrix effect was determined by comparing the signal produced by an extracted blank to the peak produced by the lower limit of quantification (LLOQ). FDA guidelines require the signal of endogenous interference be no more than 20% of the LLOQ.

### Accuracy and Precision

Accuracy and precision were determined for intra- and inter-assay variability. The degree of variation was calculated by determining the % of the interpolated concentration of QCs to the expected concentration. Accuracy (% variability of accuracy = interpolated value/expected value*100) and precision (%variation of precision = standard deviation/mean value*100) were determined at 3 concentrations (40ng/mL, 400ng/mL and 800ng/mL) in triplicate. Acceptable variation for accuracy and precision was set at 15% and at 20% for the lower concentrations.

## Results

### Recovery of Tizoxanide

TIZ and IS were detected using the optimised analyte specific and global parameters found in table 1. The mean recovery was 57.7% ±7.93 and 51.8% ±9.50 for mouse plasma and human plasma, respectively. Recovery from DMEM with varying percentages of serum was 45.2% ±3.84 (50% FBS), 48.1% ±4.72 (25% FBS), 50.3% ±6.54 (12% FBS), 50.1% ±0.63 (10% FBS), 42.1% ±4.20 (5% FBS), 42.8% ±4.50 (2% FBS), 50.5% ±9.80 (0% FBS), 51.0% ±2.30 (50% human plasma) and 40.5% ±4.68 (10% human plasma). While recovery varied between each matrix, mean recovery within each matrix demonstrated high reproducibility (example recoveries are shown in Figure 1).

**Table 1.** shows the optimised parameters used to detect TIZ and TIZ-D4 (IS).

| Parent (m/z) | Product (m/z) | RT (Min) | DP (Volts) | EP (Volts) | CE (Volts) | CXP (Volts) |
| --- | --- | --- | --- | --- | --- | --- |
| Tizoxanide | *216.8 | 2.45 | -160 | -10 | -18 | -23 |
| 263.9 | **113.9 | 2.45 | -160 | -10 | -28 | -11 |
| Tizoxanide-D4 (IS) | 221.0 | 2.45 | -160 | -10 | -20 | -19 |
| 267.9 |  |  |  |  |  |  |
| Global Parameters | Curtain Gas | Collision Gas | Voltage (V) | Temperature (°C) | Gas 1 | Gas 2 |
|  | 50 | Medium | -4500 | 450 | 50 | 50 |

**Figure 1.**
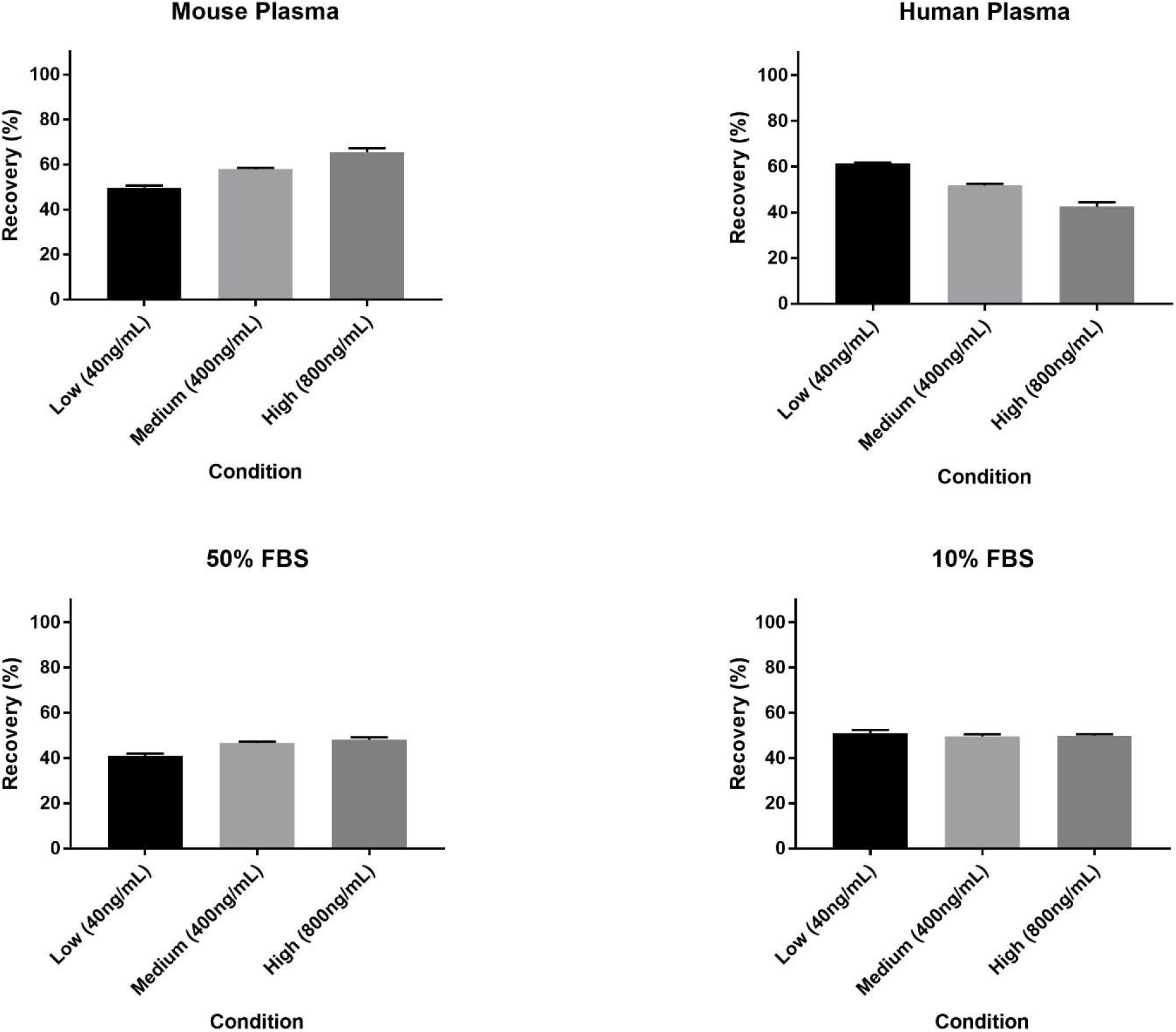
shows the recovery of TIZ in mouse plasma, human plasma, 50% FBS and 10% FBS (±SD).

### Linearity

Extracted calibrators demonstrated strong linearity (mouse plasma R^2^ = 0.9997, human plasma R^2^ = 0.9991, 50% human plasma R^2^ = 0.9983, 10% human plasma R^2^ = 0.9992, 50% FBS R^2^ = 0.9970, 25% FBS R^2^ = 0.9992, 12% FBS R^2^ = 0.9996, 10% FBS R^2^ = 0.9992, 5% FBS R^2^ = 0.9995 and 2% FBS R^2^ = 0.9995) meeting all acceptance criteria (Figure 2). The calibration curve was fit to the data using a linear equation with sample weighting of 1/X.

**Figure 2.**
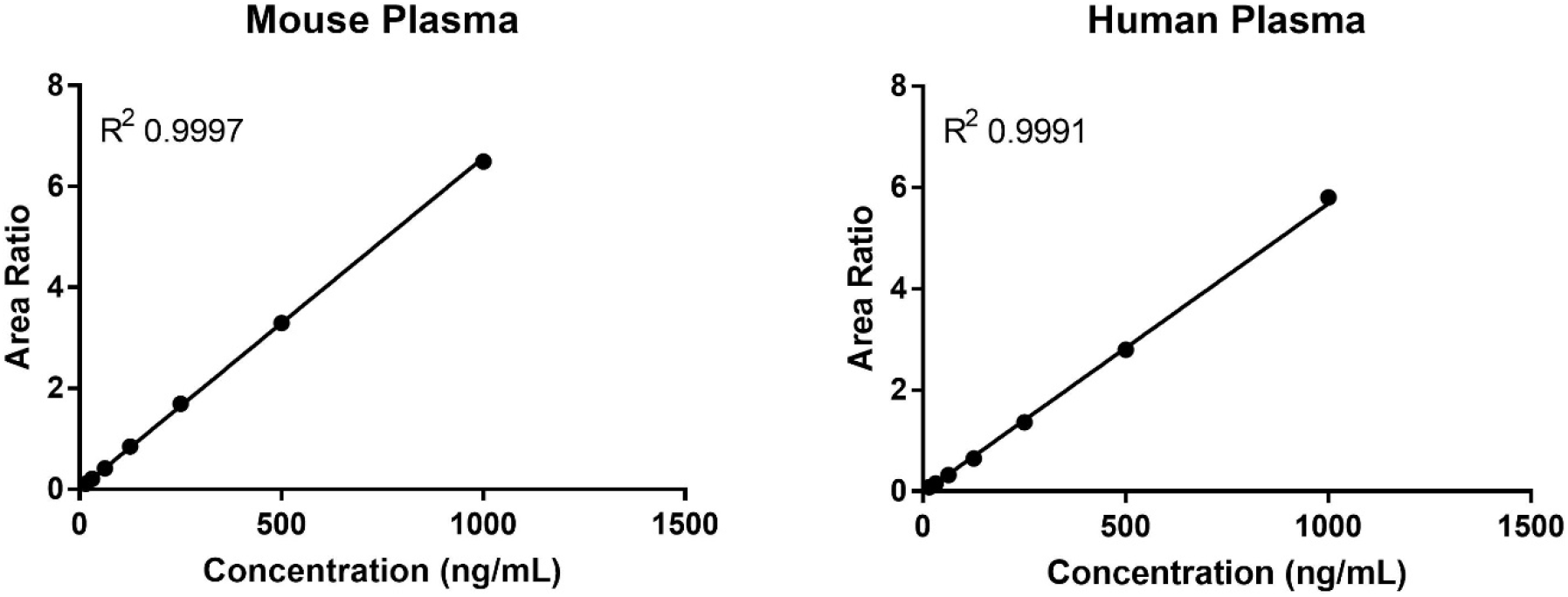
shows example standard curves produced in mouse and human plasma. Also shown is the R^2^ of the regression.

### Selectivity

Interference from endogenous compounds of a matrix is a well-documented confounding effect in bioanalysis. The Peak area produced by the LLOQ was compared to detectable signal at the retention time of TIZ (2.46 mins) and expressed as a percentage of the peak area of the LLOQ (Table 2). Representative chromatograms are shown in figure 3. Matrix effect was not deemed to interfere with the assay as endogenous signal fell below 7% in all matrices.

**Table 2.** shows the endogenous signal from an extracted blank compared to the LLOQ of TIZ and the % of the LLOQ. FDA guidelines require the signal produced by an extracted blank be no more than 20% of the LLOQ.

| **Matrix** | **Blank Signal** | **LLOQ Peak Area** | **% of LLOQ** |
| --- | --- | --- | --- |
| **Mouse plasma** | 221 | 29787 | 0.74 |
| **Human plasma** | 1291 | 88711 | 1.45 |
| **50% Human plasma** | 3134 | 99576 | 3.15 |
| **10% Human plasma** | 3071 | 75860 | 4.10 |
| **50% FBS** | 1697 | 57432 | 3.00 |
| **25% FBS** | 2683 | 58665 | 4.60 |
| **12% FBS** | 2287 | 62582 | 3.70 |
| **10% FBS** | 1641 | 85409 | 1.92 |
| **5% FBS** | 4869 | 102427 | 4.75 |
| **2% FBS** | 6571 | 100366 | 6.55 |

**Figure 3.**
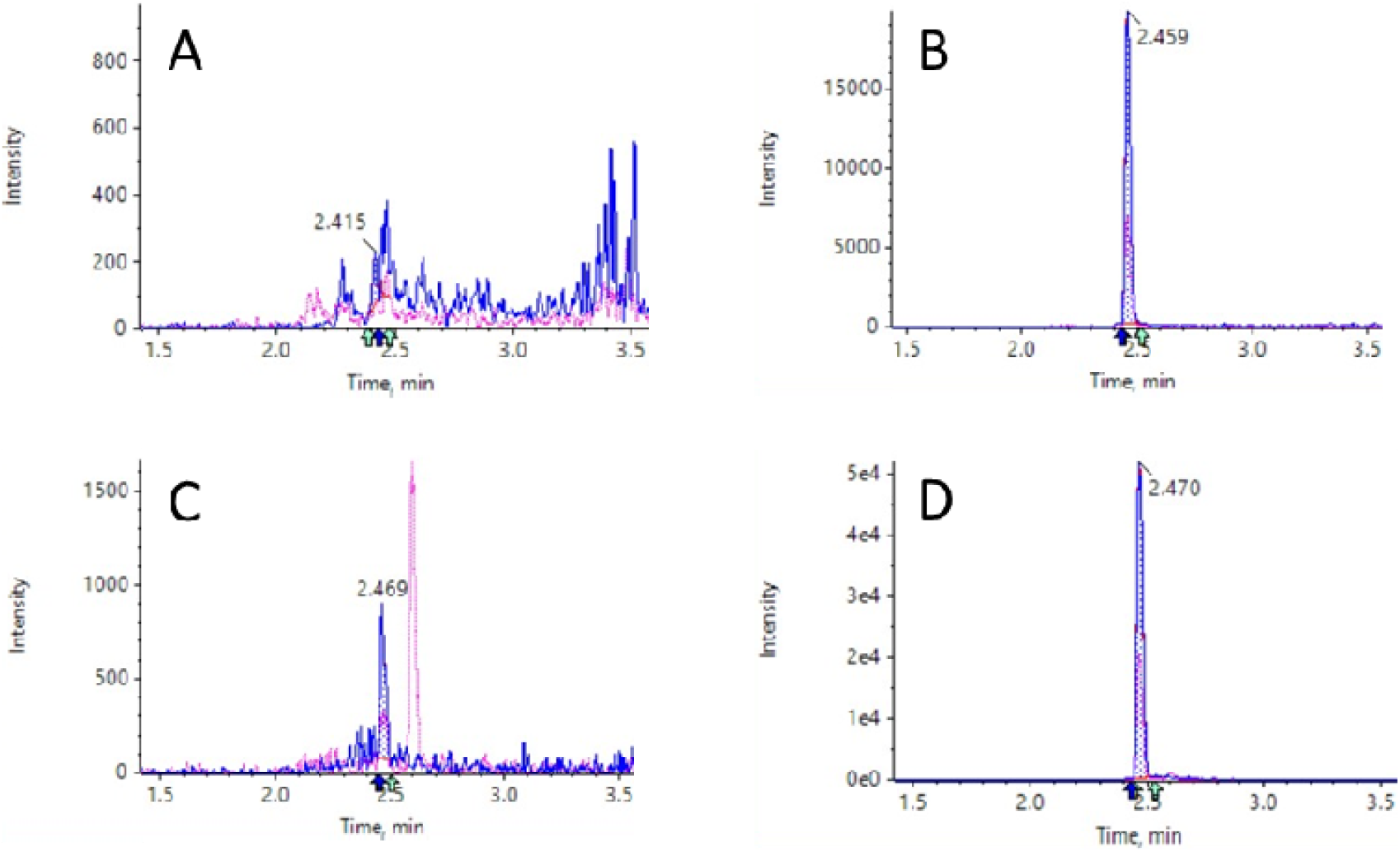
shows representative chromatograms of (A) extracted blank mouse plasma, (B) the extracted LLOQ in mouse plasma, (C) extracted blank human plasma and (D) the extracted LLOQ in human plasma.

### Accuracy and Precision

The assay described here was fully validated using mouse plasma. The variation in accuracy and precision was determined within each assay (inter-assay) and across 3 replicates of the assay (intraassay). Variance in accuracy and precision was determined by comparing the interpolated values of extracted QCs to the known values. Inter assay fell below 15% for accuracy (range −13.5% −2.2%) and precision (range 0.9% - 5.4%) at all concentrations across each replicate run. Mean intra assay variance in accuracy and precision was −3.1% and 3.6% respectively. FDA guidelines require part validation for minor changes to the assay (e.g., change of matrix). The remaining matrices were assessed for inter assay variation [28]. The intra-assay % error in accuracy was below 15% at all levels in human plasma (range −3.3 – 8.0) and FBS (range −1.4 – 10.8). Intra-assay error of precision also fell below 15% in human plasma (range 1.8 – 7.0) and FBS (range 1.1 – 7.0, Table 3).

**Table 3.** shows the validation of TIZ extracted from mouse plasma. The data shows the average quantitated sample at 3 levels, the % deviation in accuracy and the % deviation in precision. Acceptance criteria is variation no more than 15%, excluding the lower concentration where deviation may be no more than 20%.

|  |  | Intra-day |  |  | Inter-day |  |  |
| --- | --- | --- | --- | --- | --- | --- | --- |
| | | Average ± SD<br>(ng/mL) | Variance of<br>accuracy (%) | Variance of<br>precision (%) | Average ± SD<br>(ng/mL) | Variance of<br>accuracy (%) | Variance of<br>precision (%) |
| **Assay 1** | 40ng/mL | 38.5 ±1.22 | -3.8 | 3.2 | 37.0 ±2.14 | -7.4 | 5.8 |
| | 400ng/mL | 408.8 ±7.81 | 2.2 | 1.9 | 397.9 ±9.72 | -0.5 | 2.4 |
| | 800ng/mL | 798.7 ±13.9 | -0.2 | 1.7 | 789.5 ±19.36 | -1.3 | 2.5 |
| **Assay 2** | 40ng/mL | 34.6 ±1.87 | -13.5 | 5.4 | | | |
| | 400ng/mL | 390.0 ±11.29 | -2.5 | 2.9 | | | |
| | 800ng/mL | 767.3 ±38.88 | -4.1 | 5.1 | | | |
| **Assay 3** | 40ng/mL | 38.1 ±0.33 | -4.9 | 0.9 | | | |
| | 400ng/mL | 395.0 ±6.23 | 1.6 | 1.6 | | | |
| | 800ng/mL | 802.6 ±20.40 | 0.3 | 2.5 | | | |

**Table 4.**
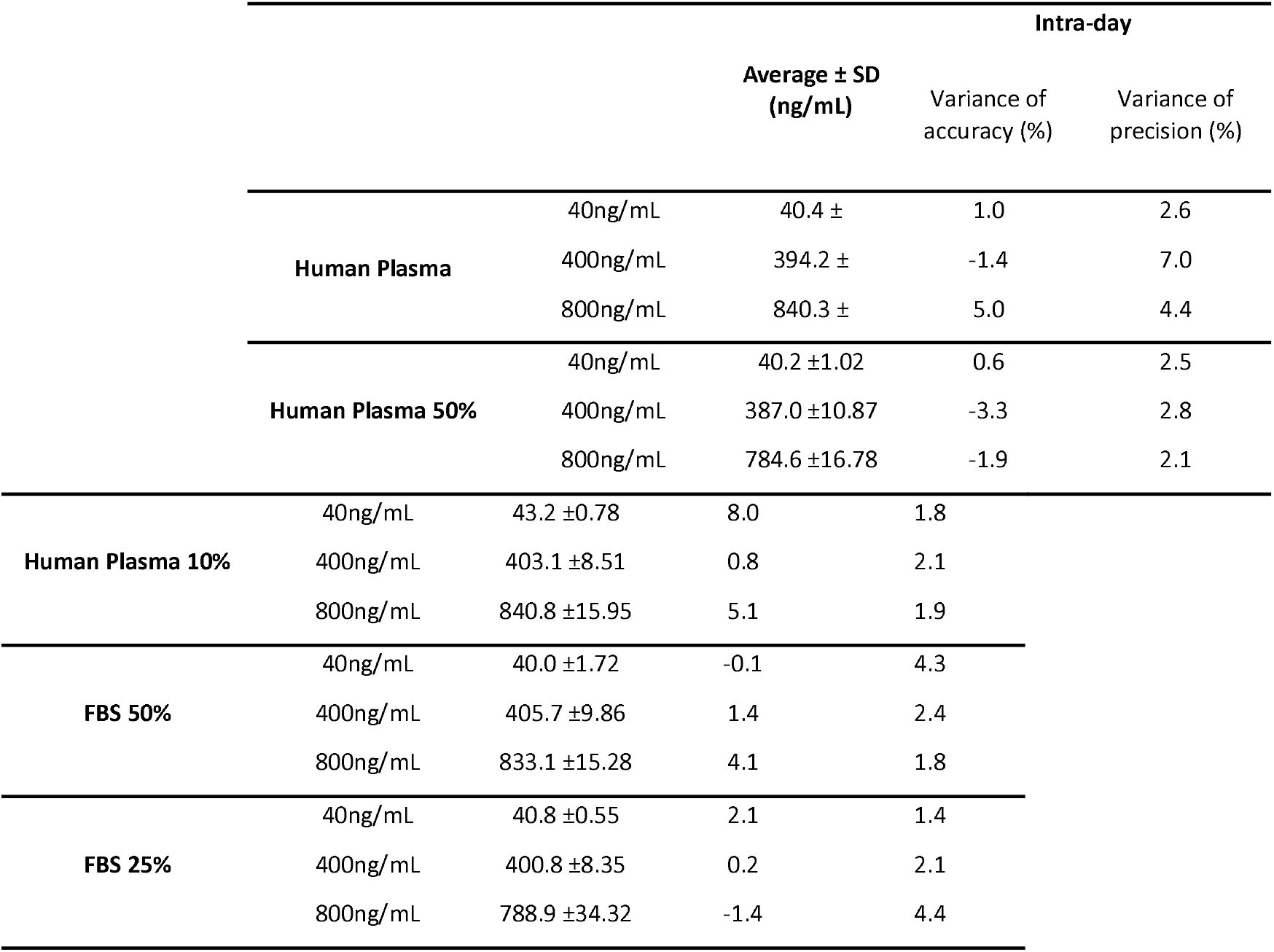
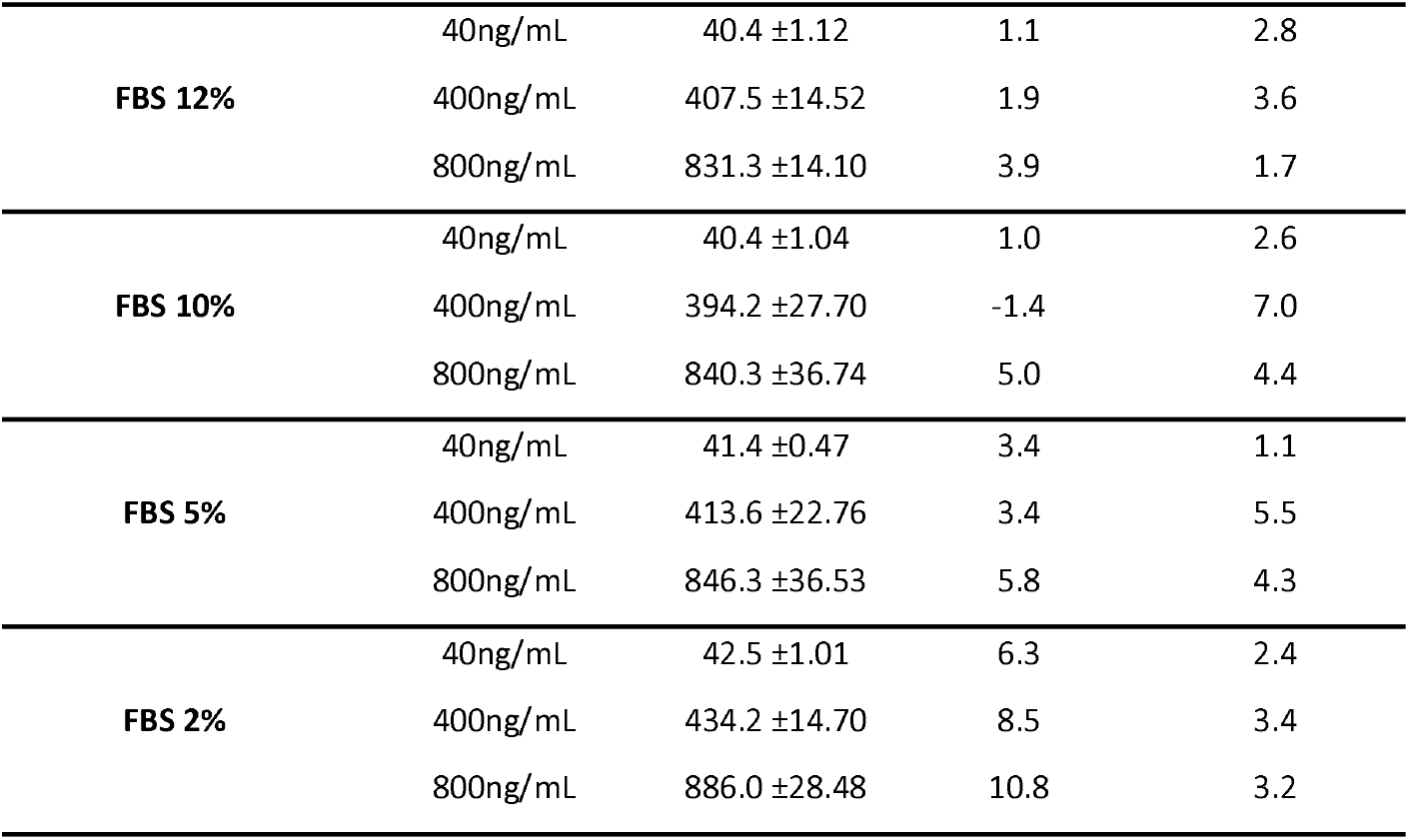
shows the part validation of each of the investigated matrices. The data shows the average quantitated sample at 3 levels, the % deviation in accuracy and the % deviation in precision. Acceptance criteria is variation no more than 15%, excluding the lower concentration where deviation may be no more than 20%.

## Discussion

The anti-SARS-CoV-2 activity of NTZ is being investigated *in vitro, in vivo*, and in clinical trials. Unlike many candidates being explored for direct repurposing as SARS-CoV-2 antivirals, the pharmacokinetics of NTZ do suggest that antiviral concentrations can be achieved in humans when benchmarked against targets derived from *in vitro* studies. However, uncertainty about high protein binding of TIZ in plasma, the specific mechanism of action, and the appropriate dose necessitates a careful understanding of the exposure-response relationship. A recent multicenter, randomised, double-blind, placebo-controlled trial, in adult patients presenting up to 3 days after onset of symptoms and with PCR-confirmed SARS-CoV-2 infections indicated some antiviral activity at 500mg three times a day [30]. However, much more data are required to support the candidacy of NTZ (including at higher doses), and studies should incorporate an understanding of the pharmacokinetics during SARS-CoV-2 infection.

A greater understanding of the pharmacokinetic-pharmacodynamic relationship requires robust, validated LC-MS/MS methods with utility in relevant biological matrices. The selection of matrices within this study included human and mouse plasma and DMEM plus 50%-2% FBS or 50%-10% human plasma. These were chosen for their relevance to clinical, *in vitro* and *in vivo* investigations of NTZ as a SARS-CoV-2 therapeutic. Although human and mouse plasma have some homogeneity, species differences can lead to changes in assay behaviour. As demonstrated by protein binding being commonly lower in preclinical species than humans [31]. Furthermore, differences in *in vitro* methods can often result in differential percentages of FBS being used to supplement DMEM in different studies. Given that previously outlined matrix effects have the potential to impact assay performance, and that this problem could be further compounded by the high protein binding observed for TIZ, >99.9% in plasma [2], a thorough investigation of matrix effect was warranted. Small differences in recovery of TIZ within different matrices were evident, but all recovery percentages were seen to be reproducible for each of the matrices (deviation <15%). Furthermore, the endogenous signal from each of the matrices was seen to be less than 7% of the LLOQ, falling well within the 20% acceptance criteria set out in FDA guidance [28]. Thus, this matrix effect is unlikely to meaningfully impact assay performance.

The decision to focus on TIZ exclusively was driven by the observation that NTZ is rapidly converted to TIZ, and that previous studies have not detected NTZ in plasma, the main matrix of interest [32]. The assay was able to quantify TIZ within multiple matrices, generating highly reproducible, precise and accurate results, indicating a tolerance to the negative impact of matrix effects. The described assay will be further developed to include the major metabolite of TIZ, tizoxanide glucuronide. In summary, a highly sensitive TIZ LC-MS/MS assay is presented, which fully meets FDA bioanalytical method development guidelines.

